# Identification and assessment of cardiolipin interactions with *E. coli* inner membrane proteins

**DOI:** 10.1101/2021.03.19.436130

**Authors:** Robin A. Corey, Wanling Song, Anna L. Duncan, T. Bertie Ansell, Mark S.P. Sansom, Phillip J. Stansfeld

## Abstract

Integral membrane proteins are localised and/or regulated by lipids present in the surrounding bilayer. Whilst bacteria such as *E. coli* have relatively simple membranes when compared to eukaryotic cells, there is ample evidence that many bacterial proteins bind to specific lipids, especially the anionic lipid cardiolipin. Here, we apply molecular dynamics simulations to assess lipid binding to 42 different *E. coli* inner membrane proteins. Our data reveals a strong asymmetry between the membrane leaflets, with a marked increase of anionic lipid binding to the inner leaflet regions of membrane proteins, particularly for cardiolipin. From our simulations we identify over 700 independent cardiolipin binding sites, allowing us to identify the molecular basis of a prototypical cardiolipin binding site, which we validate against structures of bacterial proteins bound to cardiolipin. This allows us to construct a set of metrics for defining a high affinity cardiolipin binding site on (bacterial) membrane proteins, paving the way for a heuristic approach to defining more complex protein-lipid interactions.

## Introduction

Cells are partitioned and encapsulated by biological membranes that are formed from a complex mixture of different lipids. Here, the lipids provide the necessary hydrophobic environment required to localise and tether the proteins to and/or within the membrane, acting as a solvent for the membrane-spanning region of the protein. In addition, specific interactions between particular membrane lipids and discrete regions on the surface of the protein can be of considerable importance, controlling how the protein folds, localises and functions (Corradi *et al*., 2019). Therefore the lipid composition and distribution can have a major impact on the regulation of cell membrane activity.

The identification of specific protein-lipid interactions has been tackled for a number of different proteins. In the well-studied model Gram-negative bacteria *Escherichia coli*, for instance, which has a relatively simple plasma membrane, the anionic phospholipid cardiolipin (CDL) has been shown to interact specifically with several membrane proteins, including AmtB (Patrick *et al*., 2018), SecYEG (Corey *et al*., 2018), formate dehydrogenase-N (Jormakka *et al*., 2002) and LeuT (Gupta *et al*., 2017)(Corey *et al*., 2019)(Bolla *et al*., 2020). However, there has been little in the way of systematically modelling CDL interactions with a range of different bacterial proteins in a single study.

Protein-lipid interactions are frequently studied using computational methods, such as with molecular dynamics (MD) simulations (Corey, Stansfeld and Sansom, 2020)(Corradi *et al*., 2019).These allow analysis of a given protein-lipid interaction with a high spatial and temporal resolution, as well as allowing a relatively unambiguous assignment of molecular species. In particular, use of a coarse-grained (CG) biomolecular force field, such as Martini (Siewert J Marrink *et al*., 2007; Luca Monticelli *et al*., 2008) has proven very powerful (Corradi *et al*., 2018). By reducing the degrees of freedom of a given system, sampling is improved, albeit with an associated loss in chemical resolution. This permits the dynamic modelling of protein-lipid interactions, which typically occur on the µs timescale.

Here, we use CG simulations to analyse lipid interactions with 42 *E. coli* inner membrane proteins, with each protein simulated in simple bacterial membranes. Global analysis of the data shows a strong bilayer asymmetry, with substantially more anionic lipid binding in the inner leaflet of the membrane, particularly for CDL. This is primarily driven by an increased number of lipid-facing basic residues on the cytoplasmic face of the membrane – extending the well-established positive inside rule (von Heijne, 1986) to residues which interact with the membrane. We then resolve over 700 discrete CDL binding sites from the dataset, and submit these to analyses using structural bioinformatics and free energy calculations. The data allow us to describe rules for a high affinity CDL binding site on bacterial membrane proteins. Finally, we validate our rules against previously-determined CDL sites on bacterial membrane protein structures, revealing excellent agreement.

## Results

We constructed and simulated 42 different protein/membrane systems of *E. coli* inner membrane proteins using the coarse-grained (CG) Martini force field (Siewert J. Marrink *et al*., 2007; Luca Monticelli *et al*., 2008). Symmetric membranes were built using POPE, POPG and CDL, at a 7:2:1 ratio and simulated for 5 × 5 µs: in total, we generated over 1 ms of simulation data (Figure 1B). We used this data to firstly analyse the global properties of protein-lipid interactions in the model *E. coli* membrane, and then to identify and characterise specific protein-CDL interactions.

**Figure 1.**
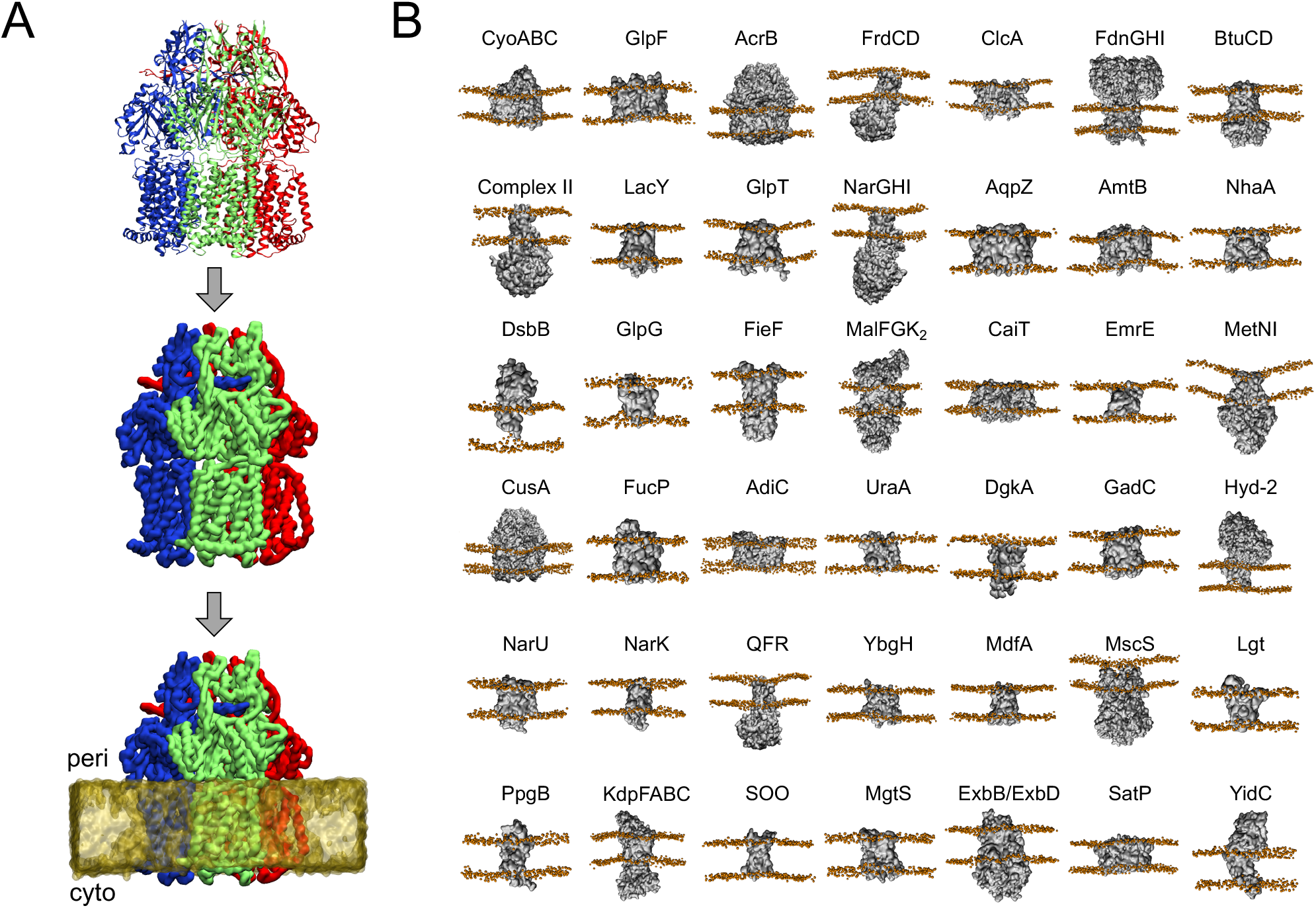
Overview of the methodology. A) Views of an example protein (AcrB, PDB 1IWG), coloured according to chain, shown in the input atomic resolution (top), in Martini CG description (middle) and embedded in a Martini CG lipid membrane (bottom). B) Views of all 42 proteins analysed in this study, with their common protein names shown above. Protein coordinates are shown in gray, phosphate beads in orange. PDB and UniProtKB IDs for each system can be found in Supplementary Table 1.

### Distribution of residues in contact with the membrane

Firstly, we carried out a global analysis of nature of protein-lipid interactions in our dataset. Across the 42 systems we see that CDL, and to a lesser extend PG, binds with a high propensity to the proteins (Figure 2A). Moreover, there is a strong asymmetry with regards to the inner (cytoplasmic) and outer leaflets, with CDL in particular far more likely to bind the protein when in the inner leaflet of the membrane. Looking at the distribution of residues in contact with the membrane (Figure 2B), this is explained by both Arg and Lys being substantially more prevalent in the inner leaflet (−2 nm) than the outer leaflet (+2 nm). Non-basic residues are evenly distributed between the two leaflets (Supplementary Figure 1B). This substantiates that the previously-asserted ‘positive-inside’ rule for membrane protein topology (von Heijne, 1986) applies not just to distribution of residues, but also of amino acid sidechains that directly interact with the membrane.

**Figure 2.**
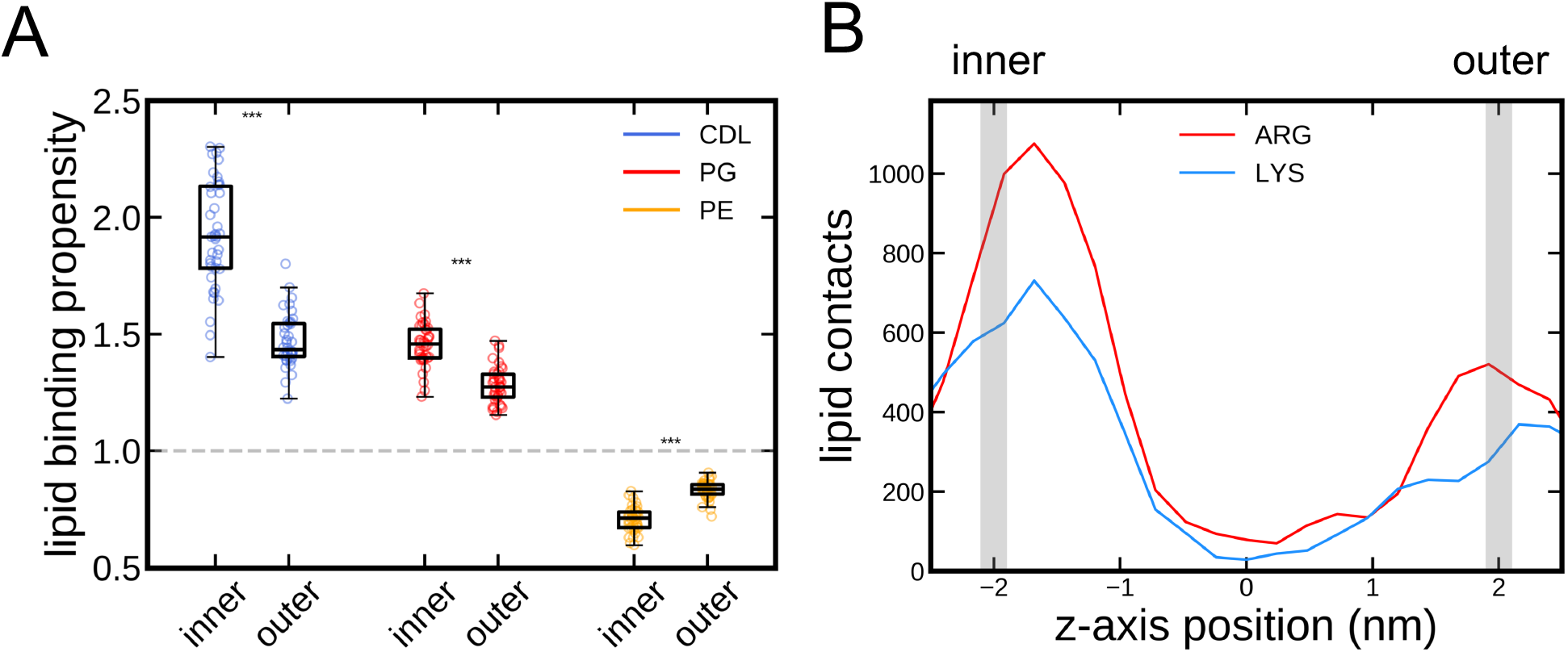
Cross-membrane asymmetry in protein-lipid interactions. A) Quantification of the number of each type of lipid in contact with the different proteins, expressed as a propensity (see Methods). Data are divided between the inner and outer leaflets, with one data point per lipid, per leaflet, per protein. Box plots show the median, upper and lower quartiles, and range (excluding flier points). Statistics are from two-tailed t-tests, with p < 0.001 in all cases. The raw data are plotted in Supplementary Figure 1A. B) Total number of Arg and Lys residues in contact with lipid molecules, plotted as a function of z-axis position, centered on the centre-of-mass of the membrane. Substantially more contacts are made in the inner leaflet than the outer. The same analysis for other residues are in Supplementary Figure 1B.

### Analysis of CDL-residue interactions

The high binding likelihood of CDL, and the seeming importance of Arg/Lys interactions in this, led us to build interaction profiles for CDL and each residue type. As expected, the CG beads representing the CDL phosphate groups are most likely to be in contact with Arg and Lys residues (Figure 3; orange), with Arg slightly more prevalent, presumably reflecting the higher propensity for Arg in membrane-facing positions (Figure 2A). This role of basic residues in CDL binding supports previous structure-based predictions (Planas-Iglesias et al., 2015).

**Figure 3.**
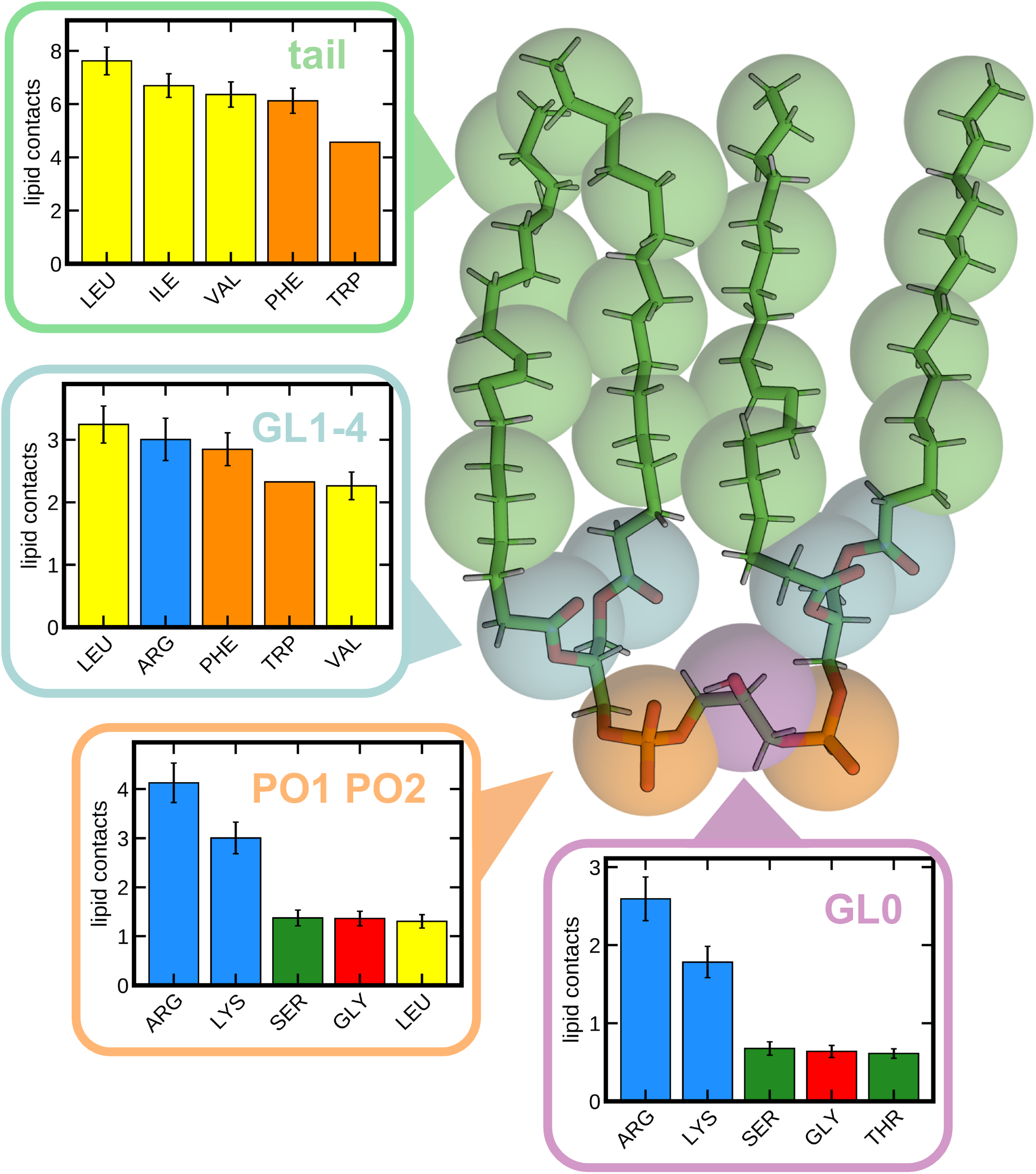
CDL-residue interaction profiles. Contact likelihood of each bead type of the Martini CDL molecule for each residue for the proteins analysed here. The 5 highest interacting residues are shown. Bar charts show mean and standard error of the mean over all 42 systems. Full residue data, and data for PE and PG, available in Supplementary Figure 2.

The central glycerol also makes substantial contacts to Arg and Lys (Figure 3; purple), with Ser, Gly and Thr residues next most likely. The similarity between the phosphate and glycerol beads is probably due to their close proximity, and the shape of the CDL headgroup. The tail-connecting glycerol beads appear to bind to aromatic residues (Phe or Trp), as well as contacting small hydrophobic residues (Leu, Val), or basic residues (Figure 3A; cyan).

### Identification of specific CDL binding sites and the importance of basic residues

We next set out to identify specific CDL binding sites from our simulation data. We followed an approach described recently (Barbera *et al*., 2018)(Duncan, Corey and Sansom, 2020), where contacts between each residue in the system and each lipid are modelled for every frame of the trajectory, and then sites are built based on residues with similar lipid-contact patterns. This analysis was run using a program designed specifically for this purpose, https://github.com/wlsong/PyLipID. From this, we identified 701 specific CDL sites with residence times above 10 ns (see Methods for filtering process). The sites had a median of 36% CDL occupancy (Figure 4A: ‘all). Representative protein structures with CDL bound are deposited at https://osf.io/gftqa/.

**Figure 4.**
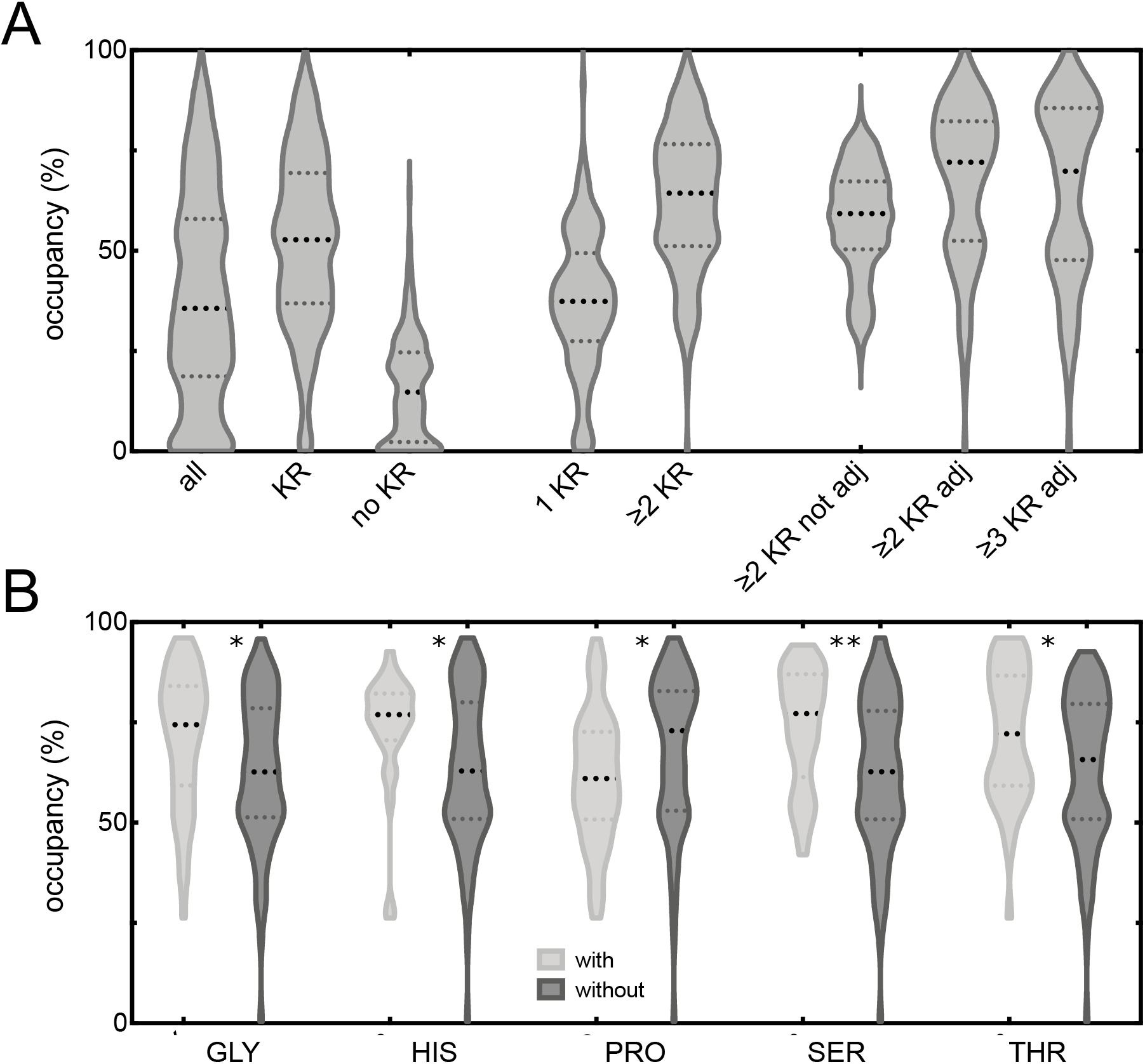
Characterisation of identified CDL sites. A) Violin plot showing the computed occupancies for identified CDL binding sites with binding durations above 10 ns. Shown are all sites (‘all’), sites with any Arg/Lys residue (‘KR’), no Arg/Lys (‘no KR’), only 1 Arg/Lys (‘1KR’), at least 2 Arg/Lys (‘≥2 KR’), and then at least either 2 or 3 structurally adjacent Arg/Lys residues (‘≥2 KR adj’ and ‘≥3 KR adj’). Reported median and interquartile range values can be found in Supplementary Table 2. C) CDL occupancies for sites with two or more structurally adjacent Arg/Lys residues and either with or without Gly, His, Pro, Ser, Thr. Statistical analysis from a two-tailed t-test, with p-values as 0.008, 0.007, 0.021, 0.001 and 0.0183, respectively.

Based on the data in Figure 3, it seems reasonable to predict that the presence of Arg or Lys residues would affect the affinity of the site. Of the 701 sites, *ca*. 60% contain at least one Arg or Lys residue, and these have a median CDL occupancy of *ca*. 53% (Figure 4A: ‘KR’), as opposed to just 14% for sites without a basic residue present (Figure 4A: ‘no KR’). We also saw a higher number of sites and median occupancy for sites with at least one basic residue in the cytoplasmic vs periplasmic leaflet (Supplementary Figure 3A).

For the 60% of sites which do contain an Arg or Lys residue, the mean number of basic residues for each site was 1.9±1.3, with an overall site size of 6 residues (Supplementary Figure 3B). As such, we looked at the impact at having two or more basic residues in the site, and saw this gives an even higher site occupancy of 64% (Figure 4A: ‘≥2KR’), as opposed to 37% for only 1 basic residue (Figure 4A: ‘1KR’).

Visualising some example sites produced by PyLipID reveals that most sites have 2-3 basic residues in very close proximity to one another (Supplementary Figure 4). Therefore, we filtered the sites based on the presence of two or more adjacent basic residues (i.e. within 0.8 nm; Supplementary Figure 3C). 32% of sites with basic residues contain adjacent Arg/Lys residues, and that these residues are typically very close on the z-axis (median 0.21 nm; Supplementary Figure 3D). Strikingly, the median occupancy of these sites (72% for ≥2 basic residues; 70% for ≥3) is far higher than sites with 2 or more Arg/Lys residues which are not adjacent (59%; Figure 4A).

Together, these observations suggest that higher occupancy CDL sites in *E. coli* membrane proteins contain 2 or more basic residues which are adjacent, i.e. within 0.8 nm, and within 0.2-0.3 nm on the z-axis.

### Other features of a two basic residue site

Of the 138 identified sites with adjacent Arg/Lys residues, we analysed other features which were associated with higher CDL binding. Firstly, analysis of the type of secondary structure the Arg/Lys residues are on reveals there is no preference for these residues to be either on helix or loop regions of the protein (Supplementary Figure 5A).

Then, we looked at other residues present in the binding sites. Several residue types appear to contribute to CDL binding likelihood, including Gly (*ca*. 36% of sites), His (*ca*. 22% of sites), Ser (*ca*. 30% of sites), and Thr (*ca*. 27% of sites), which all increase the median occupancy of the CDL site (Figure 4B). This fits well with the observation that Ser, Gly and Thr all have high levels of CDL headgroup interactions (Figure 3). Conversely, Pro (*ca*. 33% of sites), decreases the median occupancy of the CDL site.

### Contribution of different residues to the CDL poses

To assess the contributions of different residues to CDL binding, we performed alanine-scanning free energy perturbation (FEP) calculations. Here, a positive ΔΔG value indicates a higher affinity for CDL than PE (see Methods for details). We applied this approach to selected residues in 10 different binding sites, for a total of 94 mutations (see Figure 5A for three example sites). The data shows a reasonable range in the values for each residue type, with the primary observation being that Arg/Lys residues have a median interaction energy of 1.6 (0.8-2.4) kJ mol^−1^ for CDL over PE (Figure 5B), i.e. they interact with CDL more strongly than with PE. Of note, in some cases the substitution of Arg/Lys for Ala decreases the strength of the CDL interaction. This occurs in cases where there are 4 or more basic residues in total in the site, suggesting that once 2-3 basic residues are present, the addition of further basic residues diminishes the strength of the CDL coordination.

**Figure 5.**
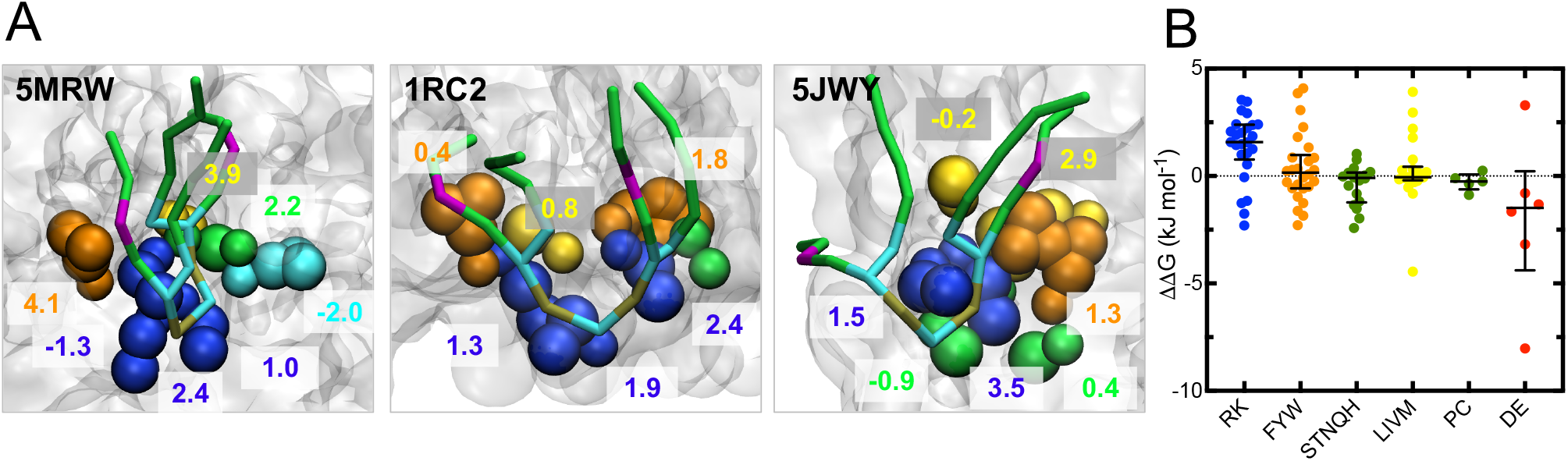
FEP Alanine Scanning. A) Views of selected CDL sites for which Ala-scanning FEP calculations were run. Each residue is shown in coloured spheres, and its calculated contribution to CDL binding reported as mean (n=10). The bound CDL from the input pose is shown as gold, cyan, green and purple sticks. B) ΔΔG values for each residue type, as calculated using FEP. Each dot is an individual residue to alanine mutation. The median and interquartile range are shown. The values for each residue are broken down in Supplementary Figure 6C and the full data in Supplementary Table 3.

In addition, certain aromatic residues show a preference for CDL over PE (Figure 5B), supporting the prediction in Figure 3. However the median interaction energy is close to, 0 so these residues need to be assessed with respect to the overall composition of the site.

### CDL binding site rules and experimental validation

Taking the data together, it is clear that a high affinity CDL site from our data set has a few key features; namely 2-3 adjacent basic residues in the same plane of the membrane, one or more polar residues, and an aromatic residue slightly further into the membrane. To evaluate these rules using experimental data, we analysed structures previously deposited in the PDB. Firstly, a direct comparison of our data with the bound CDL in *E. coli* formate dehydrogenase-N (PDB 1KQF (Jormakka *et al*., 2002)) reveals that our CG model correctly predicts the structural site, with a very high (74±24 %) CDL occupancy across the subunits (Figure 6; ‘1KQF’), and that the site follows the rules outlined above.

**Figure 6.**
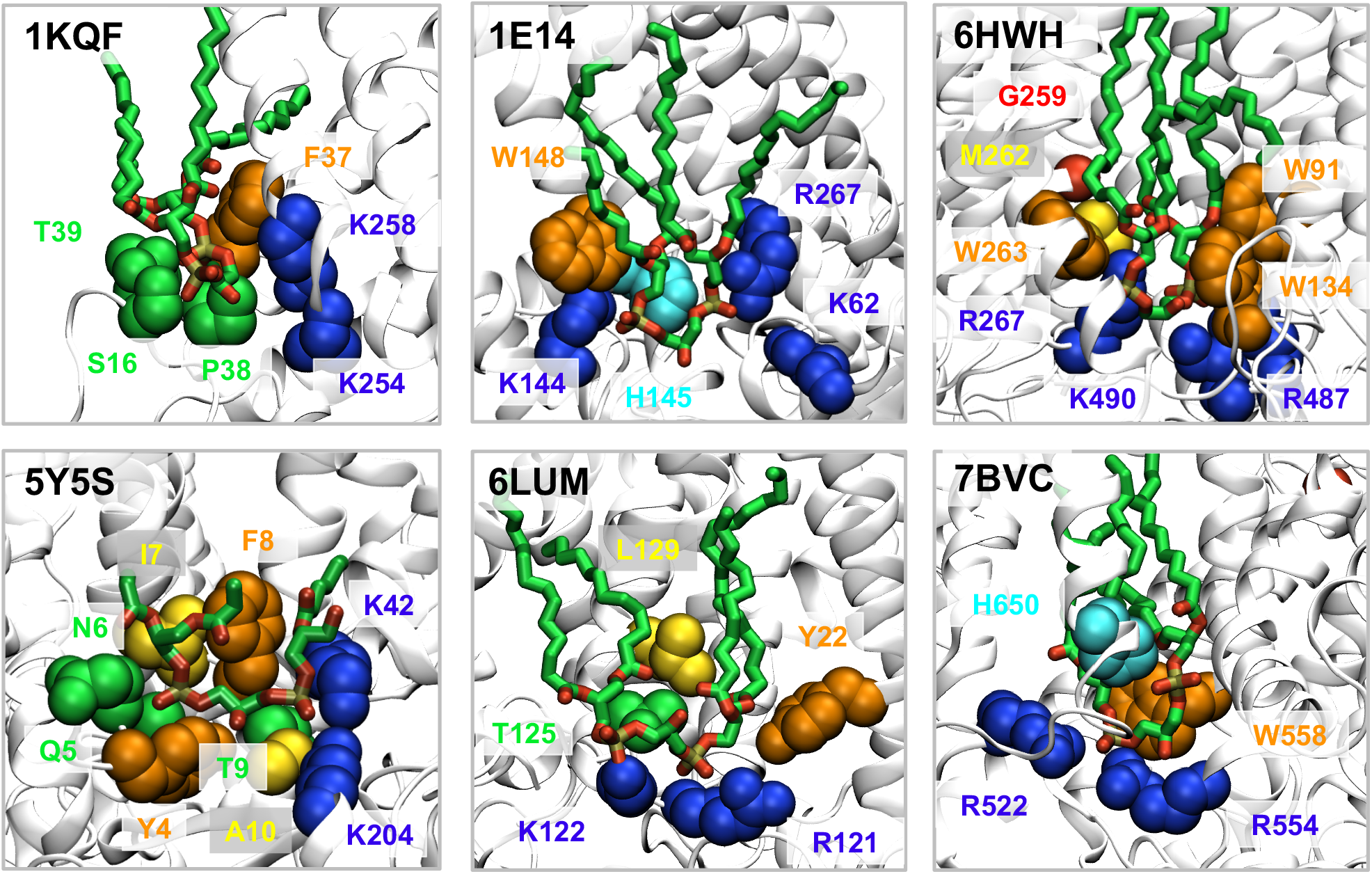
CDL sites from structural studies. Views of CDL-binding sites as determined via structural analyses, coloured as in Figure 5. See Supplementary Table 4 for a summary of all 18 identified sites with respect to the proposed CDL-binding rules.

We then carried out a broader analysis, identifying a further 17 CDL sites across 5 additional proteins (from (Fyfe *et al*., 2000) (Gong *et al*., 2020) (Wiseman *et al*., 2018) (Zhang *et al*., 2020) (Yu *et al*., 2018); see Methods for details). We compared these to our CDL site rules, observing excellent agreement (Figure 6 and Supplementary Table 4).

## Discussion

Membrane proteins bind to, and are often regulated by, many different lipids from the surrounding membrane. A number of studies have attempted to detect and probe these interactions, usually focusing on one system at a time, with notable exceptions (Corradi *et al*., 2018). Here, we investigate interactions between membrane proteins and lipids in the bacterial inner membrane, focusing on systems for which high resolution structural data of the *E. coli* membrane protein exist.

Our analyses reveal a striking pattern of asymmetry between the inner and outer leaflets of the membrane, with anionic CDL and PG binding much more readily to the inner leaflet region of the protein. This might impact the ratios of lipids in each leaflet of the membrane – if CDL and PG are sequestered at high affinity binding sites on proteins they are plausibly more likely to avoid recycling, as seen for mitochondrial CDL (Xu *et al*., 2016), contributing to a net asymmetry between the leaflets of the membrane. It is unclear what the biological necessity for this is: the proposed stabilisation of membrane proteins (Ghosh *et al*., 2020) or role as a proton sink (Haines and Dencher, 2002) in mitochondrial inner membranes could easily act in both leaflets of the membrane. In addition, the asymmetry could be in place to help balance the charges arising from the positive inside rule (von Heijne, 1986). Experimental analyses looking at CDL distribution in the membrane, like those similarly performed for PE distribution (Bogdanov *et al*., 2020), would be useful to confirm these findings.

In addition, our data reveals a set of rules for a high affinity CDL binding site on an *E. coli* – and therefore likely bacterial – membrane protein. These are:

1. 2-3 basic residues in close proximity – i.e. within 0.8 nm of each other, within 0.2-0.3 nm of each other on the z-axis, and roughly 1.8 nm from the centre of the membrane. These likely coordinate the two phosphates of the CDL molecule (Figure 3). FEP analyses suggest that each basic residue will contribute on average 1.6 kJ mol^−1^ to CDL binding above that of PE – and sometimes up to 4-5 kJ mol^−1^, and suggests that more than 3 basic residues is not necessary or desirable for a CDL site.
2. The presence of at least one polar residue – e.g. Gly, Ser, Thr or His. These are often in a similar plane to the basic residues, and are possibly important for stabilising the CDL headgroup, particularly the central glycerol.
3. One or more aromatic residues, slightly further into the plane of the membrane. These probably coordinate the glycerol groups connecting the phosphate headgroup to the acyl tails.

CDL is also highly abundant and functionally important in the mitochondrial membrane, where it has been shown to bind specifically to a wide range of proteins, including Tim23/Tim50 (Malhotra *et al*., 2017), F-ATPase (Duncan, Robinson and Walker, 2016) and Complex I (Jussupow, Di Luca and Kaila, 2019). It would therefore be interesting to extend these analyses to mitochondrial proteins, to see how universal our proposed CDL-binding rules are.

Certain caution should be drawn from the use here of a CG model of CDL. This reduces the accuracy with which interactions are defined, particularly in terms of electrostatic interactions and structural changes within the protein, but is currently essential to permit sufficient sampling for proper statistical analyses. Considerable success has been achieved using CG to model protein-lipid interactions (Corradi *et al*., 2019)(Corey, Stansfeld and Sansom, 2020), however future work incorporating fully atomistic data will permit additional insight into the data presented here, and allow a higher degree of accuracy when distinguishing between similar sites with different lipid binding properties. The increased chemical resolution will also likely make data interpretation more difficult, necessitating the use of more advanced statistical analyses.

Our study focuses principally on lipid headgroups, with little analysis of the contribution of lipid tails to binding. As a necessary simplification we chose to use simple palmitoyl-oleyl tails, where oleyl was chosen to represent the bacterial vaccenyl tail group. Therefore future analyses might also be important to investigate lipid tail diversity (Pluhackova and Horner, 2021).

## Methods

### Building systems

We referred to the MemProtMD database (Stansfeld *et al*., 2015; Newport, Sansom and Stansfeld, 2018; http://memprotmd.bioch.ox.ac.uk) to identify 42 unique *E. coli* inner membrane proteins with structural information available in the Protein Data Bank. For each protein, a single representative PDB was chosen, with the full list of PDBs used found in Supplementary Table 1. For each PDB, the atomic coordinates of the protein, embedded in a model DPPC membrane, was downloaded from the MemProtMD database. The protein coordinates were extracted, and converted to the Martini 3 open beta package v3.0.b.3.2 (Siewert J Marrink *et al*., 2007; L Monticelli *et al*., 2008). The proteins were then built into symmetric *E. coli* inner membranes using the insane protocol (Wassenaar *et al*., 2015) with 67% 1-palmitoyl-2-oleoyl-sn-glycero-3-phosphoethanolamine (POPE), 23% 1-palmitoyl-2-oleoyl-sn-glycero-3-phosphoglycerol (POPG) and 10% cardiolipin (CDL; using a −2 charge model with 23 CG beads, see Supplementary information for the topology used) in each leaflet. Note that, due to the coarse-grained nature of the Martini force field, here oleyl has been chosen to represent to common bacterial vaccenyl tail group.

Systems were solubilised with Martini 3 waters and ions to a neutral charge. Systems were minimised using the steepest descent method, then equilibrated in two rounds using 5 fs time steps for 1 ns then 20 fs time steps for 100 ns. Both equilibration steps used a semi-isotropic Berendsen barostat (Berendsen *et al*., 1984) at 1 bar, and a velocity-rescaling thermostat (Bussi, Donadio and Parrinello, 2007) at 323 K. Production simulations were then run using the Parrinello-Rahman barostat (Parrinello and Rahman, 1981) at 1 bar using 20 fs timesteps over 5 µs, running 5 repeats. All simulations were run using Gromacs 2019 (Berendsen, van der Spoel and van Drunen, 1995; Van Der Spoel *et al*., 2005).

The systems were analysed using gmx tools and MDanalysis (Gowers *et al*., 2016). Images were made with VMD (Humphrey, Dalke and Schulten, 1996) and plots made with Matplotlib (Hunter, 2007) and Prism 8.

### Modelling asymmetry in lipid contacts

For each of the 42 protein systems, the total number of each lipid type in contact with the protein was determined for both inner and outer leaflets of the membrane, as determined using the topology information present in the Orientations of Proteins in Membranes (OPM) database (Lomize *et al*., 2012). For each of the 5 repeats, average contacts (based on the distance between any protein reside and any bead from the lipid molecule being less than 0.6 nm) was taken for 0.5-5 µs of each simulation, using the Gromacs tool gmx select. Data for CDL, PG and PE binding to each protein were combined and plotted as lipid binding propensity, where propensity Is defined as:

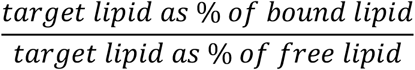

So if 20% of the lipid bound to the protein surface was CDL, and 10% of the total lipid was CDL, then the propensity would be the ratio of these, i.e. 20/10 = 2. The raw data are plotted in Supplementary Figure 1A.

### Position of lipid-contacting residues across the membrane

As the vast majority of the systems had planar bilayers, to establish a profile for protein-lipid interactions across the span of the membrane simulations were aligned according to the lipid phosphate beads, such that the centre of the membrane was set to 0 nm on the z-axis. The probability that each residue in the system contacts any lipid over the 5 × 5 µs of data was then calculated based on a 0.6 nm cut-off. For every residue with a lipid contact probability greater than 10% of the simulation time, we extracted the z-axis position from the final frame of the PO4 bead normalised simulation. We then plotted a histogram of these residues along the z-axis.

### CDL-residue interactions

Predictions of CDL binding sites were made based on the frequency of contact of each CDL particle with different protein residues across all 42 systems. Contact was determined as the number of frames of the simulation where the specified particles from the lipid and residue were within 0.6 nm, calculated using MDAnalysis. For each bead type, the 5 highest contacting residue types were plotted, with all residues plotted in Supplementary Figure 2 for all three lipid types.

### Identification of lipid binding sites

Identification of CDL binding sites was performed following a kinetic analysis of residue-lipid interactions, based on (Barbera *et al*., 2018) and (Duncan, Corey and Sansom, 2020). The program we wrote for this purpose is available at https://github.com/wlsong/PyLipID, with full details to be published separately. In brief, our approach determines whether each possible lipid/residue pair is in contact at each frame of the simulation, and then uses graph theory to cluster residues with high likelihood of simultaneously binding the same CDL headgroup. For this a double cut-off model is used: once the lipid-residue distance is smaller than a first cut-off of 0.55 nm, it is considered bound until the distance goes over a distance of 1 nm. A dual cut-off is used to account for variability in the lipid position within the binding site due to random fluctuations. Only CDL was analysed, and only interactions involving the 3 head group beads (GL0, PO1, PO2).

For each site, a global occupancy of the site was calculated based off the number of frames that CDL spends in contact with at least one residue in the site (frames_bound_):

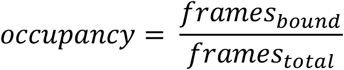

From the 42 systems we identified 986 CDL binding sites, with binding site residence times (the time the lipid is continuously in contact with any residue from the site) ranging from extremely short (*ca*. 1 ns time scale) to 2-3 µs in length. To simplify our data, we chose only sites the 701 sites with calculated binding site residence times above 10 ns, as any time below this threshold is likely just accounted for by random diffusion of the CDL molecule. The inclusion of these sites doesn’t affect the main outcomes of the study (e.g. Supplementary Figure 3E).

In addition, any individual residues with binding occupancies below 10% of the total site occupancy were removed before analysis.

### Identification of adjacent basic residues in sites

To determine which sites contain adjacent basic residues, we extracted the Cartesian coordinates of the BB bead of all Arg or Lys residues from the identified site from the input model. If an individual site has two or more basic residues within 0.8 nm in 3D space, we classified these as an adjacent pair. 0.8 nm was chosen as a reasonable cut-off based off the distribution of the distance between basic residues in sites with 2 or more basic residues present (Supplementary Figure 3C).

### Alanine scanning FEP

For identified sites with CDL occupancies above 50%, as determined using PyLipID, alanine scanning free energy perturbation (FEP) calculations were performed on any residue in contact with the CDL for at least 50% of the overall site occupancy, apart from Gly and Ala (being too similar to Ala). For this, the selected residues were alchemically perturbed to Ala through conversion of their sidechains SC beads to dummy particles with no LJ or Coulombic interactions. We ran these FEP calculations in in the presence or absence of CDL to measure the effect of mutation on lipid binding (as per (Pipatpolkai *et al*., 2020))

Poses for each site comprising the protein and bound CDL were produced using the PyLipID program. These were then embedded into a solvated Martini POPE membrane, using the insane protocol. The systems were minimized using steepest descents, and equilibrated for 10 ns using 20 fs time steps, as described above. The lipid was kept in the binding site using a 1,000 kJ mol^−1^ nm^−2^ flat bottom restraint between the COM of the CDL headgroup and the COM of the site residues, applied using plumed 2.2.3 (Bonomi *et al*., 2009; Tribello *et al*., 2014). For calculations of the system without CDL, the CDL molecule was deleted, and a 100 ns equilibration simulation was run to allow the membrane to equilibrate around the protein.

For selected residues, FEP calculations were carried out by perturbing the system with the native residue to that where the residue has dummy particles for its side chain, effectively performing a mutation to Ala. Coulombic and Lennard Jones (LJ) parameters were switched separately over the λ coordinate, over 17 windows with certain windows overlapping, following this scheme:

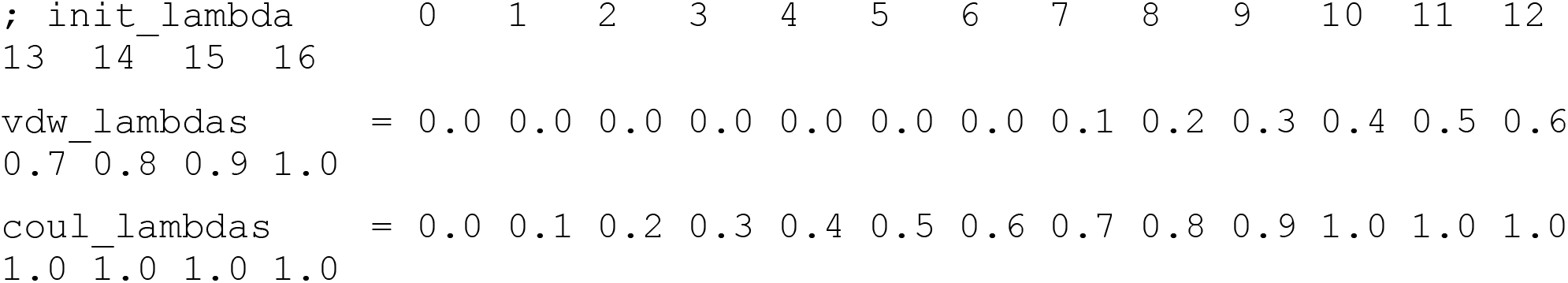

Each λ window was run for 10 repeats for 12 ns, with the first 2 ns discarded as equilibration. The separate windows were constructed into energy landscapes along λ using Multistate Bennett Acceptance Ratio (MBAR) (Shirts and Chodera, 2008) as implemented in alchemical analysis (Klimovich, Shirts and Mobley, 2015). Convergence is shown for one case (5JWY) in Supplementary Figure 6A.

ΔΔG values were computed using the cycle in Supplementary Figure 6B. Here, the determined energy cost of substituting a residue to an alanine when bound (ΔG_arg>ala.CDL_) and not bound (i.e. in a pure PE membrane; ΔG_arg>ala.PE_) to a CDL molecule. A positive ΔΔG (ΔG_arg>ala.CDL_ − ΔG_arg>ala.PE_) means that the residue is interacting more strongly with the CDL than with a generic lipid.

### Analysis of PDBs

The PDB was queried for the Chemical ID “CDL”, giving 222 structures (as of Feb 2021), 64 of which were bacterial. Filtering out duplicate entries for the same system left 6 unique structures, with 18 CDL sites. Comparison with the proposed CDL rules were made based off visual inspection (see Supplementary table 4). Note that PDBs containing modified fluorescent CDL derivatives were not included in this analysis.

## Acknowledgements

This work was supported by the Wellcome [208361/Z/17/Z], the BBSRC [BB/P01948X/1, BB/R002517/1 BB/S003339/1] and MRC [MR/S009213/1]. This work used ARCHER UK National Supercomputing Service (http://www.archer.ac.uk) and JADE, provided by HECBioSim, the UK High End Computing Consortium for Biomolecular Simulation (hecbiosim.ac.uk), which is supported by the EPSRC (EP/L000253/1). We acknowledge the use of Athena at HPC Midlands+, which was funded by the EPSRC on grant EP/P020232/1, and the University of Warwick Scientific Computing Research Technology Platform for computational access.

## Supplementary Information

**Supplementary Figure 1.**
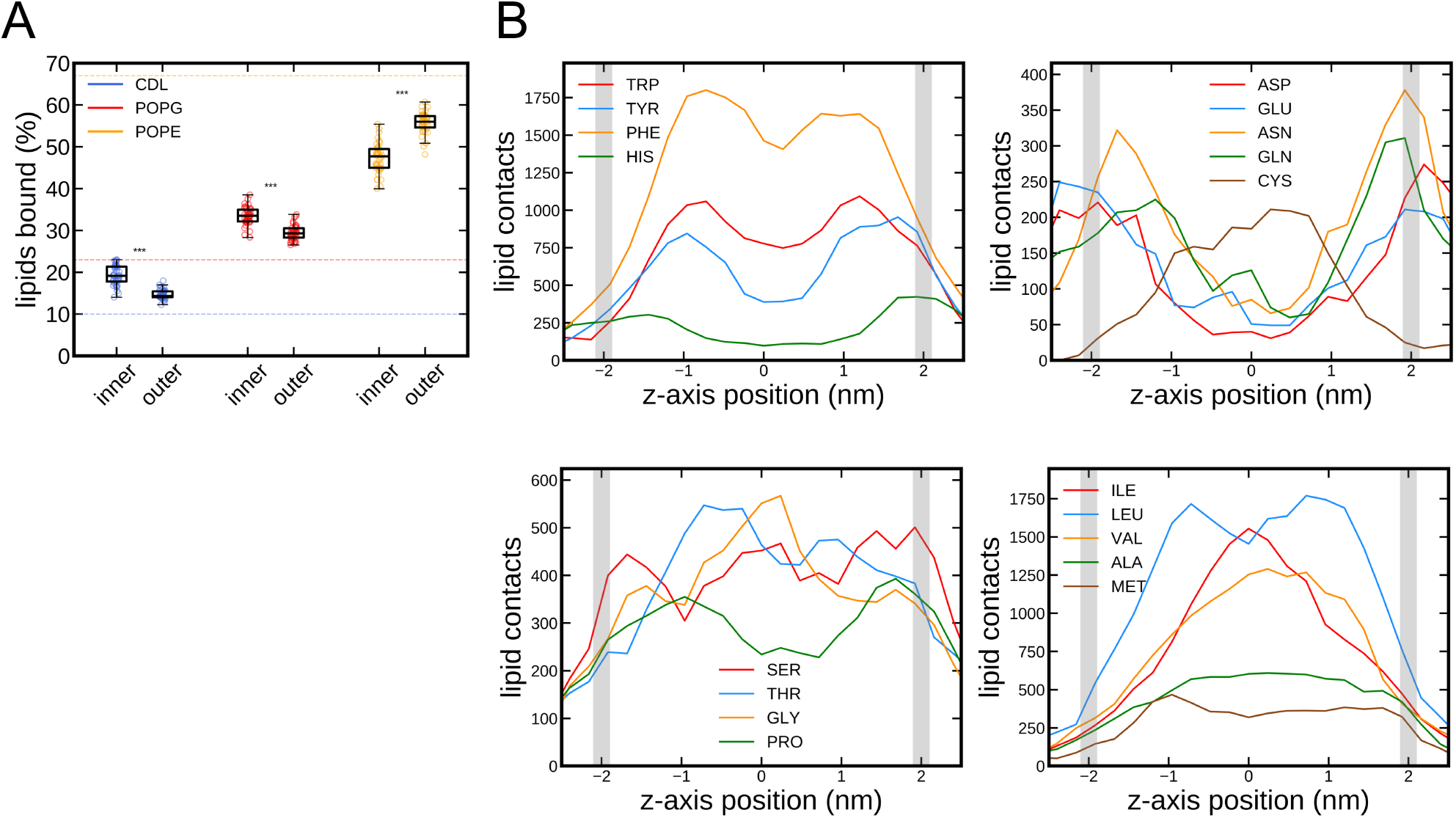
**A)** As Figure 2A, but plotting instead the raw binding probability for each lipid. Dotted lines indicate the total concentration of each lipid in the membrane. **B)** As Figure 2B, but for all residues except Arg and Lys.

**Supplementary Figure 2.**
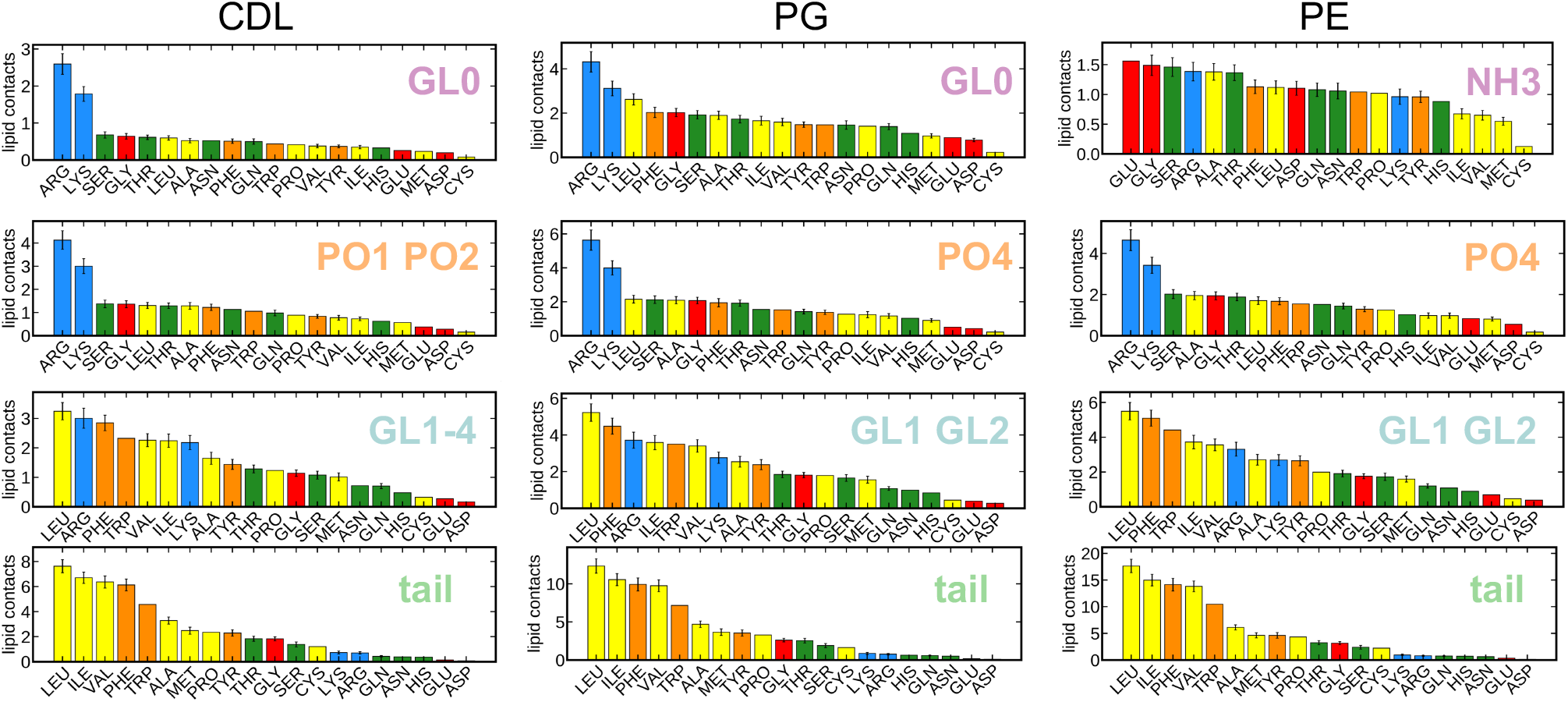
As per the data in Figure 3, but showing the binding likelihood of all residues to CDL, POPG and POPE.

**Supplementary Figure 3.**
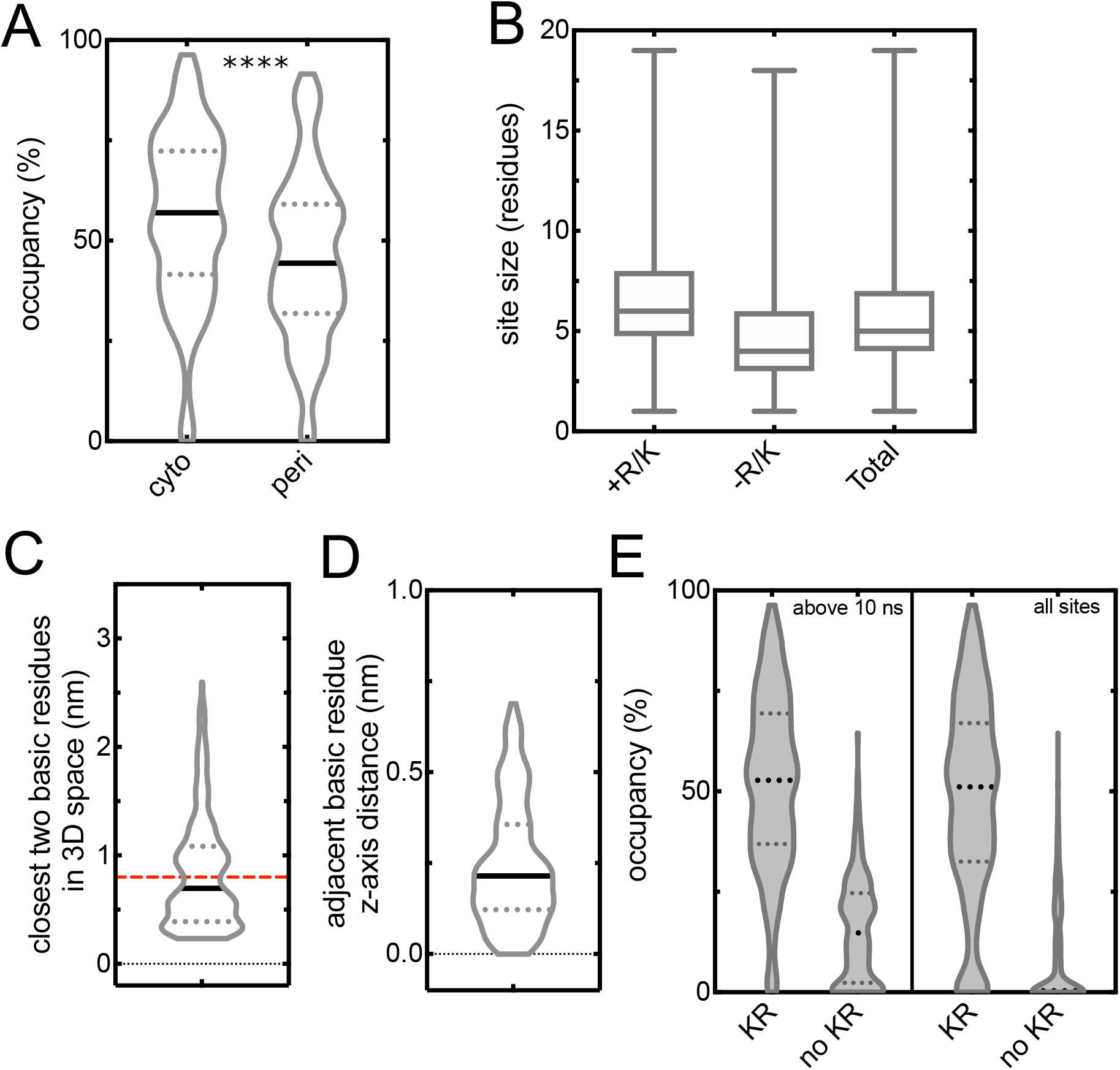
**A)** Occupancies for all sites containing at least one basic residue, separated by which leaflet the site is located. p < 0.0001 from a two-tailed t-test. **B)** Box plot showing the computed sizes of the identified sites, for sites with an Arg or Lys (‘+R/K’), without (‘−R/K’) and total. Shown are lines for the median, upper and lower quartiles, and range. **C)** Distribution of distance between closest two basic residues in the 255 sites identified with 2 or more basic residues. A red line denotes the chosen 0.8 nm cut-off. **D)** Distribution of z-distances between basic residues pairs determined to be adjacent. **E)** As Figure 4A, but also including data for sites with binding durations less than 10 ns. Note that the left two plots are the same as in Figure 4A.

**Supplementary Figure 4.**
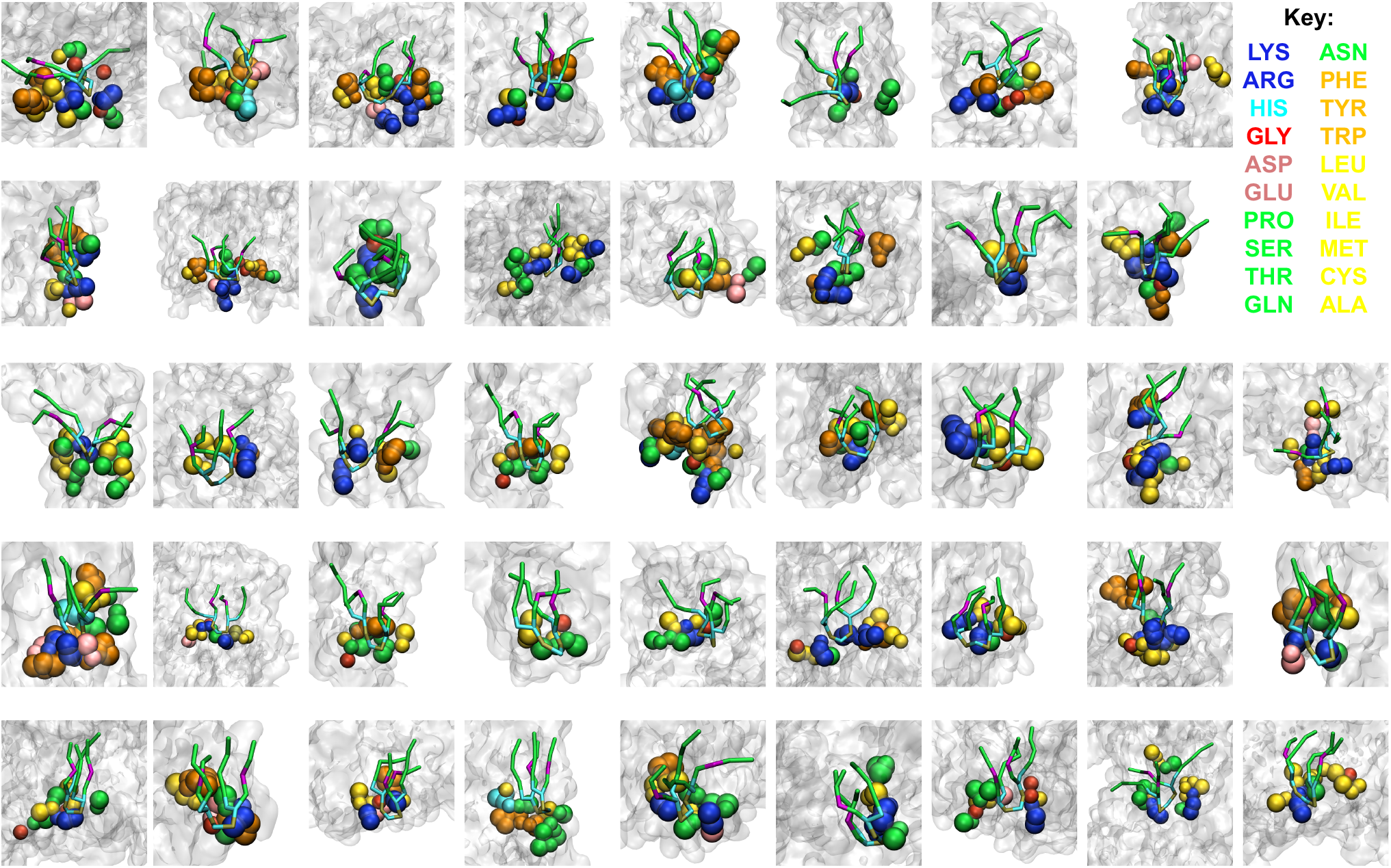
Views of selected CDL sites, as identified from PyLipID. Protein backbones are shown as gray surface, with sites residues coloured according to type. The bound CDL molecule in a representative pose is shown in coloured sticks Note that the sites are aligned with the CDL oriented with the headgroup at the bottom, whether the site is cytoplasmic or periplasmic

**Supplementary Figure 5.**
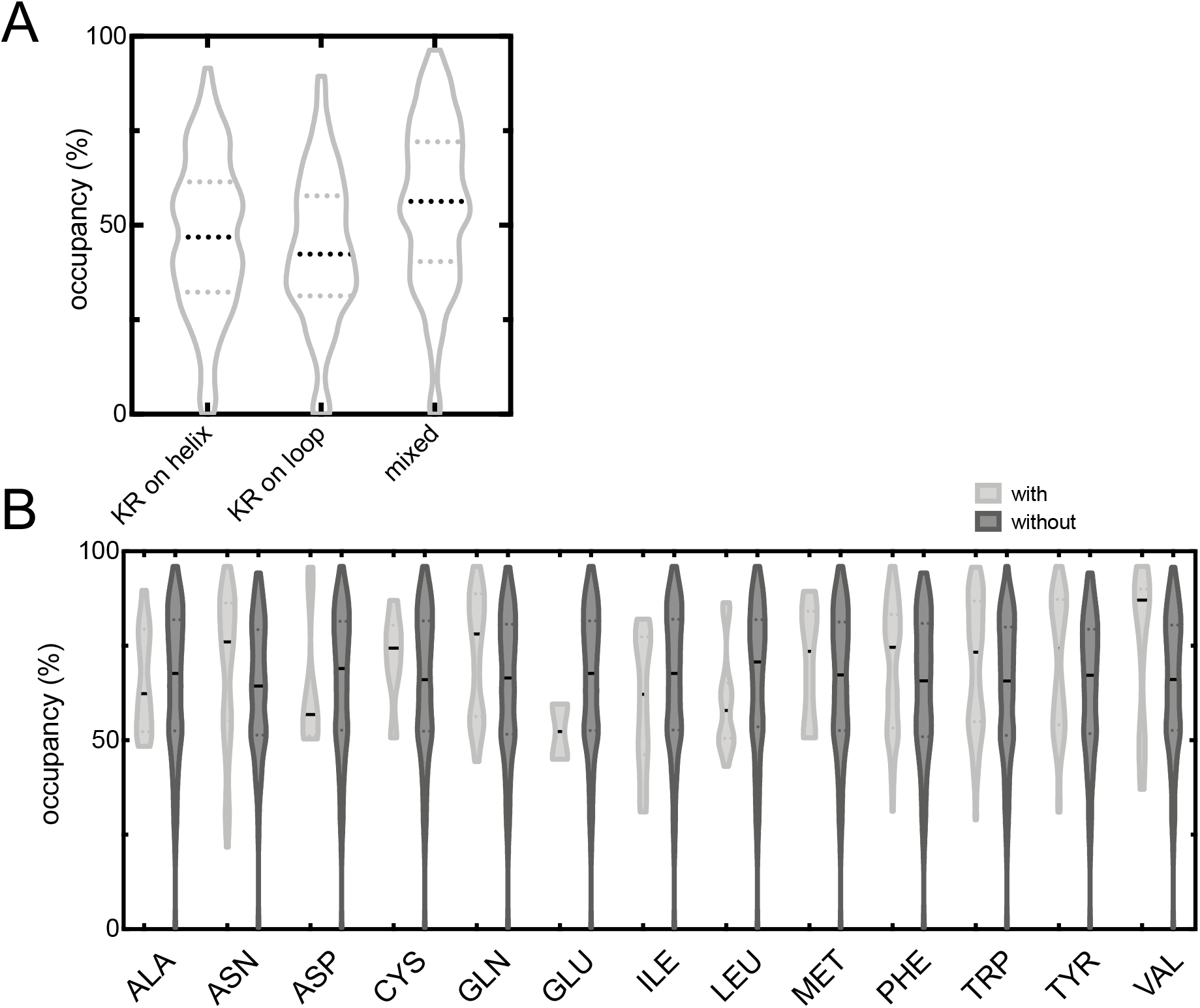
**A)** Analysis of type of structure the CDL site basic residues are found on. ‘Helix’ includes all CG beads with definition H, 1, 2 and 3, and ‘loop’ all beads with type E, T, S and C, as defined by the martinize script (https://github.com/cgmartini/martinize.py). Shown are data for sites the basic residue solely on helix (‘KR on helix’), solely on a loop (‘KR on loop’), or a combination of both (‘mixed’). **B)** As Figure 4B but for residues with no significant difference as determined using a two-tailed t-test.

**Supplementary Figure 6.**
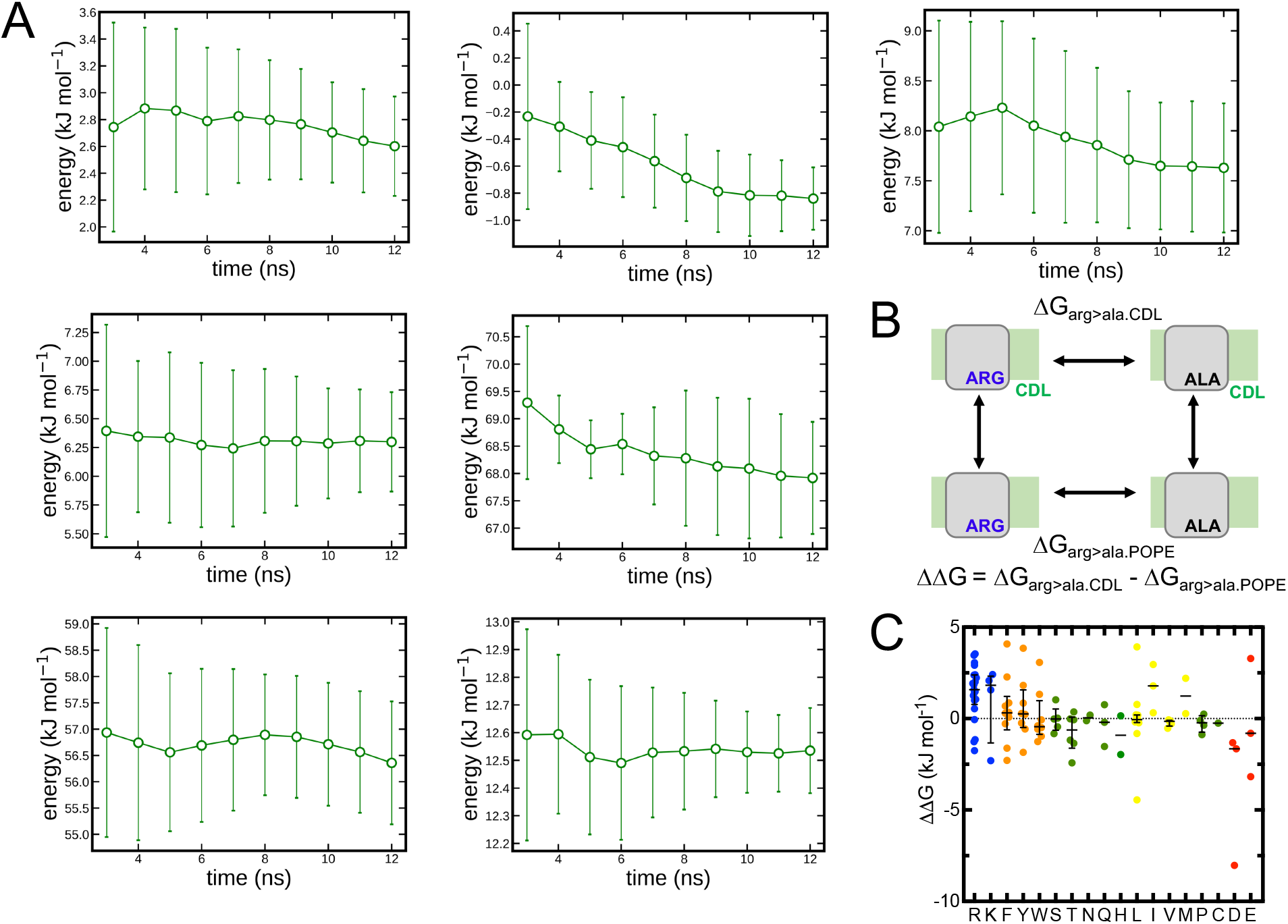
**A)** Convergence analysis of a subset of alanine scanning FEP calculations (5JWY chosen as representative site). For each residue, 2 ns is discarded as equilibration, and MBAR performed on increasing lengths of simulation up to 10 ns (12 ns in total). All calculations converge within 10 ns. Mean and standard deviation of 10 repeats plotted. **B)** Thermodynamic cycle for perturbing an arginine residue to alanine in either the presence (ΔG_arg>ala.CDL_) or absence (ΔG_arg>ala.POPE_) of a bound CDL molecule. The interaction energy between the residue and the CDL is then calculated as the difference between these values. In the schematic, the protein is gray, the membrane is green, and the residue of interest is highlighted. **C)** As Figure 5B, but for each residue separately. Shown as bars is the median and interquartile range, where more than 3 values are present.

**Supplementary table 1.** Details of protein structures used in this analysis.

| **UniProtKB ID** | **PDB ID** |
| --- | --- |
| SATP_ECOLI | 5ZUG |
| FDNH_ECOLI<br>FDNI_ECOLI | 1KQF |
| GLPG_ECOLI | 2IC8 |
| YNEM_ECOLI | 5OQT |
| NARK_ECOLI | 4JR9 |
| DSBB_ECOLI | 2HI7 |
| FUCP_ECOLI | 3O7P |
| KDGL_ECOLI | 3ZE3 |
| ATKA_ECOLI<br>ATKB_ECOLI<br>ATKC_ECOLI | 5MRW |
| C561_ECOLI | 5OC0 |
| NHAA_ECOLI | 1ZCD |
| LACY_ECOLI | 1PV6 |
| ADIC_ECOLI | 3OB6 |
| NARU_ECOLI | 4IU8 |
| FRDC_ECOLI<br>FRDD_ECOLI | 1KF6 |
| FRDD_ECOLI | 4KX6 |
| URAA_ECOLI | 3QE7 |
| CLCA_ECOLI | 1KPK |
| EXBB_ECOLI<br>EXBD_ECOLI | 5SV0 |
| AMTB_ECOLI | 1U77 |
| CYOB_ECOLI<br>CYOC_ECOLI<br>CYOA_ECOLI | 1FFT |
| MSCS_ECOLI | 5AJI |
| MDFA_ECOLI | 4ZP0 |
| LGT_ECOLI | 5AZC |
| NARI_ECOLI | 1Q16 |
| GLPF_ECOLI | 1FX8 |
| YIDC_ECOLI | 6AL2 |
| FIEF_ECOLI | 2QFI |
| METI_ECOLI | 3DHW |
| ACRB_ECOLI | 1IWG |
| CUSA_ECOLI | 3K07 |
| MBHS_ECOLI<br>CYBH_ECOLI | 4GD3 |
| PGPB_ECOLI | 5JWY |
| EMRE_ECOLI | 3B5D |
| DTPD_ECOLI | 4Q65 |
| GLPT_ECOLI | 1PW4 |
| MALG_ECOLI<br>MALF_ECOLI | 2R6G |
| GADC_ECOLI | 4DJI |
| CAIT_ECOLI | 2WSX |
| DHSC_ECOLI<br>DHSD_ECOLI | 1NEK |
| BTUC_ECOLI | 1L7V |
| AQPZ_ECOLI | 1RC2 |

**Supplementary table 2.** Number of sites, and median and interquartile range of CDL occupancies from the features mentioned in the text. Data from Figure 4 and Supplementary Figures 3 and 5.

| **Feature** | **No. of sites** | **25% Q** | **Median** | **75% Q** |
| --- | --- | --- | --- | --- |
| All | 701 | 18.69 | 35.63 | 57.95 |
| KR | 430 | 36.87 | 52.78 | 69.38 |
| no KR | 271 | 2.355 | 14.77 | 24.68 |
| 1 KR | 175 | 27.5 | 37.36 | 49.42 |
| ≥2 KR | 255 | 51.18 | 64.35 | 76.58 |
| ≥2 KR not adj | 117 | 50.36 | 59.28 | 67.28 |
| ≥2 KR adj | 138 | 52.51 | 72.05 | 82.27 |
| ≥3 KR adj | 87 | 47.67 | 69.83 | 85.6 |
| KR (incl <10 ns) | 469 | 32.54 | 51.14 | 66.97 |
| no KR (incl <10 ns) | 517 | 0.016 | 0.372 | 16.91 |
| cyto | 271 | 41.57 | 56.97 | 72.32 |
| peri | 156 | 31.86 | 44.38 | 59.1 |

**Supplementary table 3.**
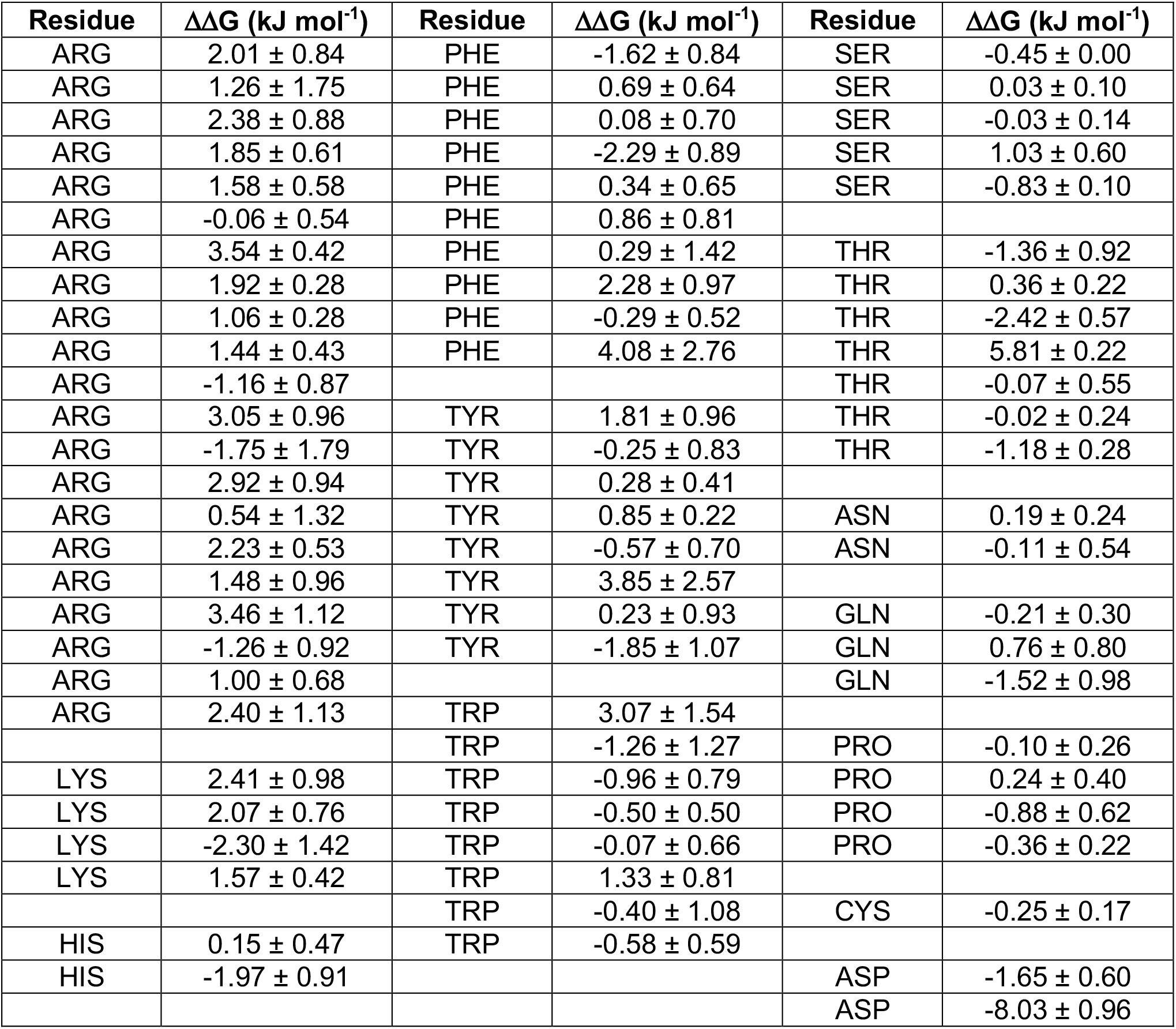
Calculated ΔΔG values from Figure 5

| Residue | $\Delta\Delta G$ (kJ mol<sup>-1</sup>) | Residue | $\Delta\Delta G$ (kJ mol<sup>-1</sup>) | Residue | $\Delta\Delta G$ (kJ mol<sup>-1</sup>) |
| --- | --- | --- | --- | --- | --- |
| ARG | 2.01 ± 0.84 | PHE | -1.62 ± 0.84 | SER | -0.45 ± 0.00 |
| ARG | 1.26 ± 1.75 | PHE | 0.69 ± 0.64 | SER | 0.03 ± 0.10 |
| ARG | 2.38 ± 0.88 | PHE | 0.08 ± 0.70 | SER | -0.03 ± 0.14 |
| ARG | 1.85 ± 0.61 | PHE | -2.29 ± 0.89 | SER | 1.03 ± 0.60 |
| ARG | 1.58 ± 0.58 | PHE | 0.34 ± 0.65 | SER | -0.83 ± 0.10 |
| ARG | -0.06 ± 0.54 | PHE | 0.86 ± 0.81 |  |  |
| ARG | 3.54 ± 0.42 | PHE | 0.29 ± 1.42 | THR | -1.36 ± 0.92 |
| ARG | 1.92 ± 0.28 | PHE | 2.28 ± 0.97 | THR | 0.36 ± 0.22 |
| ARG | 1.06 ± 0.28 | PHE | -0.29 ± 0.52 | THR | -2.42 ± 0.57 |
| ARG | 1.44 ± 0.43 | PHE | 4.08 ± 2.76 | THR | 5.81 ± 0.22 |
| ARG | -1.16 ± 0.87 |  |  | THR | -0.07 ± 0.55 |
| ARG | 3.05 ± 0.96 | TYR | 1.81 ± 0.96 | THR | -0.02 ± 0.24 |
| ARG | -1.75 ± 1.79 | TYR | -0.25 ± 0.83 | THR | -1.18 ± 0.28 |
| ARG | 2.92 ± 0.94 | TYR | 0.28 ± 0.41 |  |  |
| ARG | 0.54 ± 1.32 | TYR | 0.85 ± 0.22 | ASN | 0.19 ± 0.24 |
| ARG | 2.23 ± 0.53 | TYR | -0.57 ± 0.70 | ASN | -0.11 ± 0.54 |
| ARG | 1.48 ± 0.96 | TYR | 3.85 ± 2.57 |  |  |
| ARG | 3.46 ± 1.12 | TYR | 0.23 ± 0.93 | GLN | -0.21 ± 0.30 |
| ARG | -1.26 ± 0.92 | TYR | -1.85 ± 1.07 | GLN | 0.76 ± 0.80 |
| ARG | 1.00 ± 0.68 |  |  | GLN | -1.52 ± 0.98 |
| ARG | 2.40 ± 1.13 | TRP | 3.07 ± 1.54 |  |  |
|  |  | TRP | -1.26 ± 1.27 | PRO | -0.10 ± 0.26 |
| LYS | 2.41 ± 0.98 | TRP | -0.96 ± 0.79 | PRO | 0.24 ± 0.40 |
| LYS | 2.07 ± 0.76 | TRP | -0.50 ± 0.50 | PRO | -0.88 ± 0.62 |
| LYS | -2.30 ± 1.42 | TRP | -0.07 ± 0.66 | PRO | -0.36 ± 0.22 |
| LYS | 1.57 ± 0.42 | TRP | 1.33 ± 0.81 |  |  |
|  |  | TRP | -0.40 ± 1.08 | CYS | -0.25 ± 0.17 |
| HIS | 0.15 ± 0.47 | TRP | -0.58 ± 0.59 |  |  |
| HIS | -1.97 ± 0.91 |  |  | ASP | -1.65 ± 0.60 |
|  |  |  |  | ASP | -8.03 ± 0.96 |

**Supplementary table 4.**
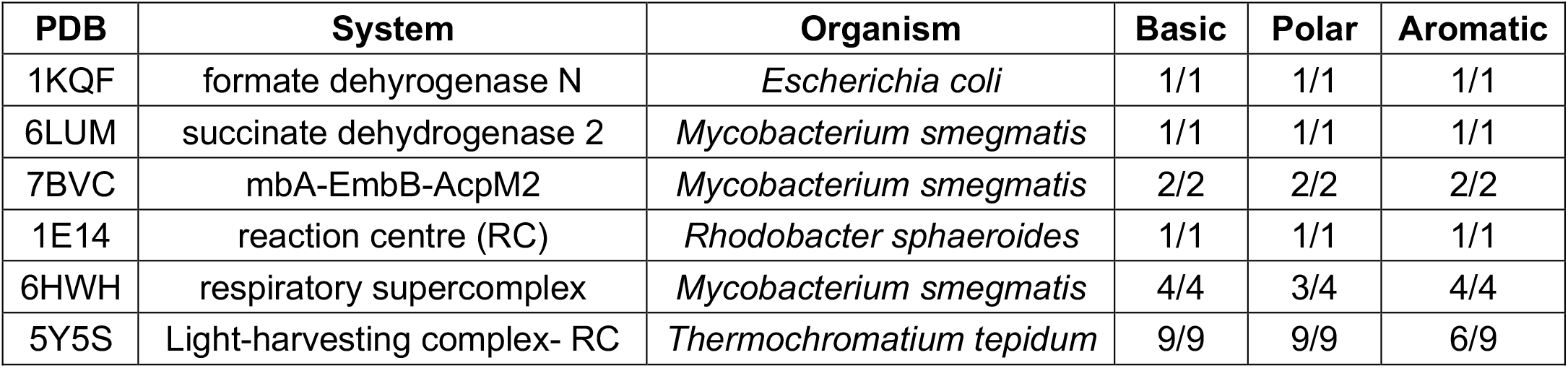
Bacterial CDL sites from the PDB, and comparison to the site outlined by our CG data in terms of ≥2 Arg/Lys residues within 0.8 nm (‘Basic’), at least one Thr/Ser/His/Gly residue (‘Polar’) and at least 1 Phe/Trp/Tyr residue further into the membrane (‘Aromatic’).

| **PDB** | **System** | **Organism** | **Basic** | **Polar** | **Aromatic** |
| --- | --- | --- | --- | --- | --- |
| 1KQF | formate dehydrogenase N | *Escherichia coli* | 1/1 | 1/1 | 1/1 |
| 6LUM | succinate dehydrogenase 2 | *Mycobacterium smegmatis* | 1/1 | 1/1 | 1/1 |
| 7BVC | mbA-EmbB-AcpM2 | *Mycobacterium smegmatis* | 2/2 | 2/2 | 2/2 |
| 1E14 | reaction centre (RC) | *Rhodobacter sphaeroides* | 1/1 | 1/1 | 1/1 |
| 6HWH | respiratory supercomplex | *Mycobacterium smegmatis* | 4/4 | 3/4 | 4/4 |
| 5Y5S | Light-harvesting complex- RC | *Thermochromatium tepidum* | 9/9 | 9/9 | 6/9 |

